# Monitoring the effects of chemical stimuli on live cells with metasurface-enhanced infrared reflection spectroscopy

**DOI:** 10.1101/2021.06.30.450584

**Authors:** Steven H. Huang, Jiaruo Li, Zhiyuan Fan, Robert Delgado, Gennady Shvets

## Abstract

Infrared spectroscopy has found wide applications in the analysis of biological materials. A more recent development is the use of engineered nanostructures – plasmonic metasurfaces – as substrates for metasurface-enhanced infrared reflection spectroscopy (MEIRS). Here, we demonstrate that strong field enhancement from plasmonic metasurfaces enables the use of MEIRS as a highly informative analytic technique for real-time monitoring of cells. By exposing live cells cultured on a plasmonic metasurface to chemical compounds, we show that MEIRS can be used as a label-free phenotypic assay for detecting multiple cellular responses to external stimuli: changes in cell morphology, adhesion, lipid composition of the cellular membrane, as well as intracellular signaling. Using a focal plane array detection system, we show that MEIRS also enables spectro-chemical imaging at the single-cell level. The described metasurface-based all-optical sensor opens the way to a scalable, high-throughput spectroscopic assay for live cells.

## Introduction

Infrared (IR) spectroscopy is a powerful technique that enables the label-free, non-destructive analysis of biomolecules through their molecular vibrational fingerprints. Although the IR absorption cross section of a typical molecule is orders of magnitude larger than that of Raman scattering, IR absorption is still relatively weak, requiring long signal collection time when conventional Fourier transform infrared (FTIR) spectrometers are used. Fortunately, analogously to the well-known surface-enhanced Raman scattering (SERS) (*1*–*3*), IR absorption can also be enhanced by several orders of magnitude through localized hotspots of electromagnetic field in the vicinity of plasmonic particles, in a process known as surface-enhanced infrared absorption (SEIRA) (*4*–*8*). Unlike SERS, the mid-IR (MIR) light used in FTIR spectroscopy has a larger (2 µm – 10 µm) wavelength and can be significantly enhanced by designer SEIRA substrates fabricated using top-down lithographic techniques. Periodic arrays of nano-antennas, nano-slits, and more complex structures, known as metasurfaces, have been engineered to have tailored optical response over a broad range of frequencies (*9*–*16*). Such plasmonic metasurfaces make excellent biosensors and have been successfully applied to the spectroscopic analysis of biomolecules including protein monolayers, lipid bilayers, fixed and dried cells, as well as cells captured through dielectrophoresis (*17*–*23*).

A rapidly expanding area of application for IR spectroscopy is the spectroscopic analysis of biological samples, such as tissue sections, cells, and serum. IR spectroscopy requires no additional preparation or staining of biological samples because their constitutive molecules – proteins, lipids, carbohydrates, and nucleic acids – have distinct vibrational fingerprints in the mid-IR spectral region that serve as endogenous labels (*24, 25*). IR spectroscopy has found application in cytological and histological diagnosis, for example in classifying cells as normal or cancerous, or classifying tissue sections into different subtypes (*26*–*32*). Remarkably, IR spectroscopy can sometimes even outperform conventional screening due to the richness of the biochemical information contained in the spectra (*33*). Typically, IR spectroscopy generates a large amount of high-dimensional spectral information that can be analyzed for feature extraction and classification using multivariate statistical analysis and machine learning. Assisted by these chemometric techniques, IR spectroscopy has been used to analyze and classify cellular responses to different biochemical compounds according to their mode of action (*34*–*37*).

Although metasurface-based SEIRA has been applied successfully to the spectroscopy of biomolecules, so far metasurfaces have not been used for the spectroscopy of live cells cultured directly on the metasurfaces – a key step that would greatly enhance the application of SEIRA. In this work, we propose and demonstrate the use of metasurface-enhanced infrared reflection spectroscopy (MEIRS) (*23*) as a live cell monitoring technique. The spectra collected from live cells are much more complex and information-rich than those from biomolecules (*18*) because cells are highly heterogeneous objects that interact with the nano-topography of the metasurface in a complex manner (*38, 39*). From this complexity flows the opportunity: developing a new highly-sensitive phenotypic cellular assay for monitoring subtle changes in cellular adhesion, cytoskeletal reorganization, and membrane composition. Such cellular assay technology would combine the precision of molecular fingerprinting derived from IR spectroscopy with the high-throughput nature of phenotypic cellular assays (*40*–*42*).

Currently, most of the standard transmission-mode FTIR micro-spectroscopy setups are unsuitable for measuring live cells because MIR light is strongly attenuated in water. Extending IR spectroscopy to live-cells typically require special setup configurations, such as thin (around 10 µm) flow cells (*43, 44*) or attenuated total reflection (ATR) setups (*45, 46*), but neither of these are suitable for scaling to high-throughput analysis. With metasurface, the absorption spectrum of the target analyte (e.g., a confluent cell monolayer cultured atop of a metasurface) is encoded into the reflectance spectrum. Thus, when fabricated on a MIR-transparent CaF_2_ substrate, metasurfaces enable collecting the absorbance spectra of live cells in the reflectance mode, which is crucial to avoiding water absorption. Metasurfaces can also be fabricated on large planar substrates compatible with modern sample handling technologies such as microplates or microfluidics, enabling the scale-up to larger, high-throughput assays.

MEIRS shares some similarity with existing optical phenotypic cellular assays based on resonant waveguide grating (RWG) (*47*–*49*) and surface plasmon resonance (SPR) (*50*–*53*) sensors in terms of device geometry and sensing modality. In these assays, cells are also seeded on engineered surfaces supporting surface or guided waves, and the cell-penetrating evanescent fields are used to probe the local refractive index of the cells. Cellular response is detected by measuring the refractive index modulation induced by evolving cell-substrate adhesion or cytoskeletal reorganization, known as dynamic mass redistribution (DMR). Cells undergo DMR in response to various stimuli, making these sensors versatile platforms for cellular assays. In particular, DMR has been shown to be an early indication of phenotypical response of cells in G-protein coupled receptor signaling pathway, with important application in drug screening (*42, 54*). Comparing to these existing phenotypical assays, MEIRS information content is qualitatively richer because, in addition to probing local refractive index change, it provides spectro-chemical information about the analyte: cellular proteins and lipids, complex culture medium and extracellular matrix, and, potentially, drug uptake and retention. This additional spectro-chemical information is instrumental in detecting changes not reflected in refractive index alone, as well as in interpreting the underlying mechanisms of complex phenotypical cellular responses.

Here, we demonstrate the application of MEIRS as a cellular assay technique by monitoring live cells responding to different chemical stimuli through the change in their IR spectra. Plasmonic metasurfaces are designed to enhance the IR absorption from cells, with particular focus on the amide I and II peaks around 1500 cm^-1^ – 1700 cm^-1^ (common to proteins) as well as CH_2_/CH_3_ vibrations around 2800 cm^-1^ – 3000 cm^-1^ (common to lipids). Spectral shifts of the metasurface resonances are also observed in the collected IR spectra, and are used to track refractive index changes around the metasurface due to adherence variations of the cells. Cells are grown on gold metasurfaces fabricated on a planar MIR-transparent CaF_2_ substrate, integrated with a flow cell. The time-varying stimulus-specific spectral fingerprint of the cell – its phenotypic response barcode (PRB) – is probed through near-field extending roughly 100 nm into the matrix behind the metasurface (*22*). The resultant signal predominantly reflects changes in cellular adhesion to the metasurface and, uniquely to MEIRS, in the cellular membrane properties. As two examples of stimulus-driven adhesion- and membrane-related cellular changes, we use MEIRS to monitor cellular signaling and subsequent detachment from a metasurface by trypsin, as well as cholesterol depletion from cellular membrane by methyl-β-cyclodextrin (MβCD).

## Results

### Fano-resonant metasurface for enhanced IR spectroscopy of cells

Finite-bandwidth metasurface resonances were matched to the wavenumbers corresponding to the molecular vibrations of interest. The important classes of biomolecules in cells include carbohydrates, nucleic acids, proteins, and lipids. CH_2_/CH_3_ peaks in the 2850 – 2950 cm^-1^ region is attributed to lipids, including intracellular lipid droplets as well as the cellular membrane. Amide I peak at *ω*_*A*1_ ≈ 1650 cm^-1^ and amide II peak at *ω*_*A*2_ ≈ 1550 cm^-1^are attributed to a combination of C=O stretching, N-H bending, and C-N stretching in the amide backbone of the proteins (*55*). Considering the localized nature of the metasurface enhancement, we are mainly targeting the CH_2_/CH_3_ and amides absorption lines; these compounds are abundant at the basal side of the cell because they originate from the cellular membrane, focal adhesion complex, and the cytoskeleton.

The metasurface geometry used in this work is adapted from those previously used for sensing protein monolayers (*18*) and fixed/dried cells (*22*). SEM images of the metasurface are shown in Fig. 1A, and its detailed geometrical dimensions are presented in Fig. S1. The metasurface supports two resonances: a super-radiant dipolar mode at *ω*_*D*_ ≈ 2900 cm^-1^, and a sub-radiant quadrupolar mode at *ω*_*Q*_ ≈ 2080 cm^-1^. The coupling between these two modes results in a Fano-type lineshape, the sharp slope of which is useful for monitoring small shifts in the resonance mode (*18*). The amide I and II bands are located on the red side of the quadrupolar mode, while the dipolar mode is matched to the CH_2_/CH_3_ peaks. In addition, the resonance shift of the quadrupolar mode is used to track refractive index modulation related to cellular changes.

**Fig. 1.**
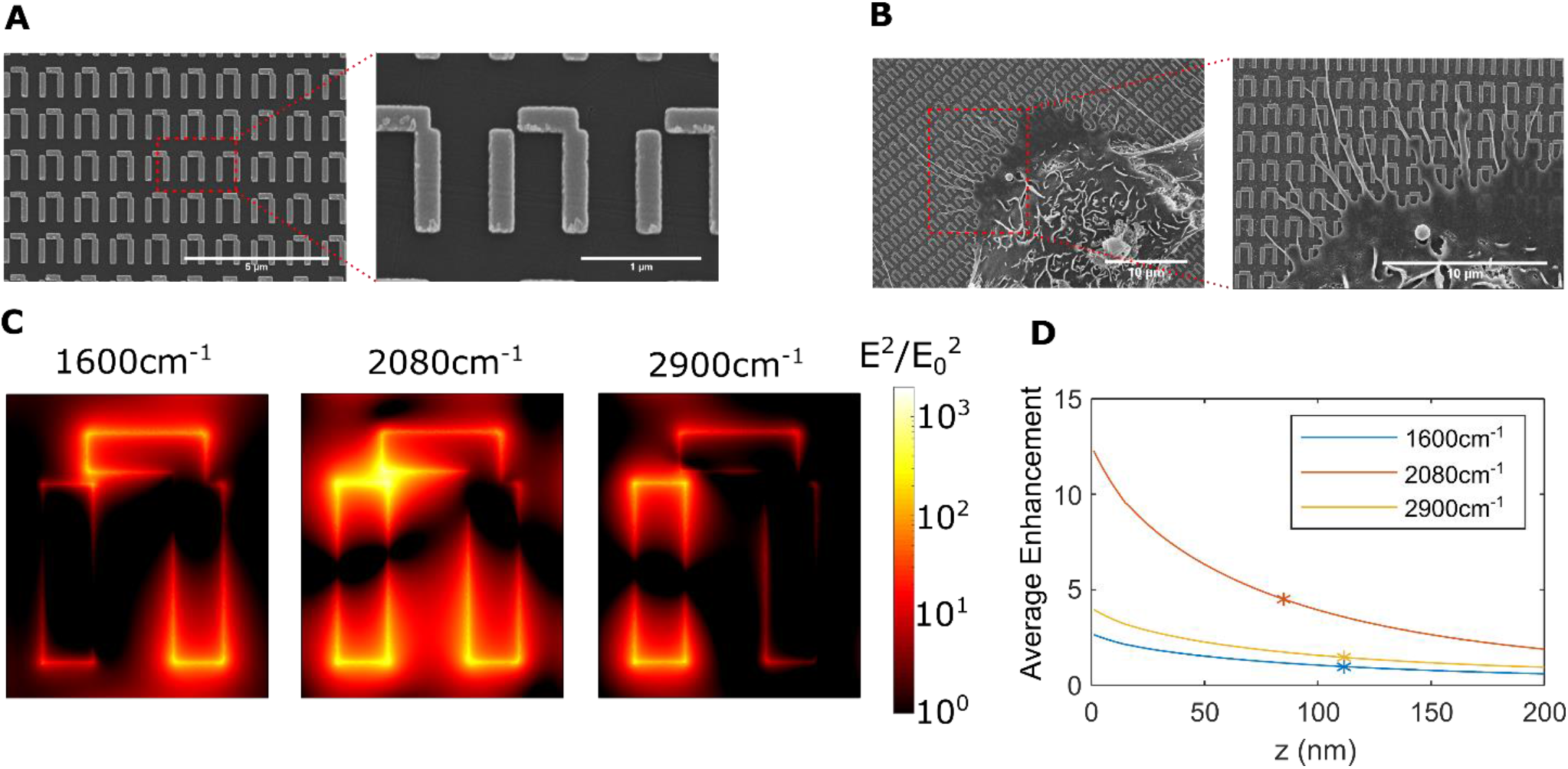
Fano-resonant plasmonic metasurface for IR spectroscopy of live cells. (**A**) SEM images of the nanostructured metasurface and (**B**) A431 cells growing on it. Note preferential attachment of filopodia to the gold nanoantennas. (**C**) Simulated near field intensity distribution at a plane *z* = 10 nm above the metasurface and (**D**) its penetration into water, at wavenumbers corresponding to amides (1600 cm^-1^) and lipids (2900 cm^-1^) absorption peaks, as well as plasmonic resonance (2080 cm^-1^). Stars: intensity reduction by 1/*e*.

The spatial distribution of the local optical field intensity *I*(*x, y, z*) differs at different wavenumbers, as shown in Figs. 1C,D. The shorter antenna on the left is mainly responsible for the resonance in the lipids absorption window (*ω* ≈ 2900 *cm*^-1^), while the antenna on the right, together with the horizontal coupler, is responsible for the resonance in the amides window (*ω* ≈ 1600 *cm*^-1^). At the Fano resonance (*ω* ≈ 2080 *cm*^-1^), both antennas interact to create a field distribution maximized at the narrow gap between the two antennas. The enhancement factor η =*I*(*x, y, z*)/*I*_0_, defined as the ratio of the local and incident optical intensities, is the highest at the Fano resonance, where *η*_max_ ∼ 10^3^ locally and ⟨*η*⟩_*x,y*_ > 10 just above the metasurface when averaged across the x-y plane. Depending on the wavenumber, the 1/e penetration depth *l*_pen_ is in the 80 nm < *l*_pen_ < 120 nm range in the vertical *z* direction. Because the thickness of an adherent cell is typically on the order of several micrometers (*56*), the metasurface does not probe the entire cell. Instead, this shallow field penetration makes the metasurface particularly sensitive to cell adhesion, cytoskeletal reorganization, as well as modulations in cellular membrane.

Because of the highly nonuniform spatial distribution of the plasmonic nearfield, the interaction between cells and the optical fields is dependent on the details of cellular attachment to the metasurface. Unlike protein monolayers and lipid bilayers that have been measured using metasurface previously (*17, 18, 20, 57*), cells are highly non-uniform objects with complex morphology and adhesion patterns that are dependent on the specific cell type under study. For this study we have focused on A431 human squamous cell carcinoma cells as a model system. Scanning electron microscopy (SEM) images of fixed A431 cells on metasurface show that the cells form filopodia that preferentially attached to the gold nanoantennas, rather than the CaF_2_ substrate (Fig. 1B). While only the focal adhesions at the periphery of the cell are accessible to SEM, it is reasonable to assume that focal adhesions beneath the cell are also preferentially formed on gold surfaces. Such adhesion pattern is beneficial for MEIRS because it increases the overlap between cell adhesion sites and the optical field localized in the vicinity of the gold antennas. This also suggests that the MEIRS signal is strongly weighted by the contribution from these adhesion sites.

MEIRS of cells cultured on the metasurface was carried out using an FTIR-coupled IR microscope. The metasurface was probed from the CaF_2_ side, with the metasurface side immersed in cell culture media as shown in Fig. 2F. In order to keep the live cells under physiological conditions during the measurement, the metasurface was attached to a polydimethylsiloxane (PDMS) flow cell maintained at 37°C, and was continuously perfused with Leibovitz’s L15 medium (unless otherwise indicated) using a programmable syringe pump. Typical reflectance spectra, with and without A431 cells on the metasurface, are presented in Fig. 2A. The Lorentzian dipolar resonance and Fano-shaped quadrupolar resonance can be seen from the reflectance spectra. The dip around 1650 cm^-1^ comes from the IR absorption of water. The IR absorption of molecules measured using MEIRS appears as dips and peaks overlapping the reflectance spectrum of the metasurface itself, originating from the coupling between the molecular vibrations and metasurface resonance (*18*). Due to the plasmonic near-field enhancement, larger vibrational peak intensities can be obtained with metasurface in comparison with undecorated substrate/cell interface (Fig. 2A). Comparing the spectra with and without the cells on metasurface, we observe small but clear spectral differences in three regions, corresponding to the two amide peaks attributed to proteins (Fig. 2B), Fano resonance shift (Fig. 2C), and multiple CH_2_/CH_3_ peaks attributed to lipids (Fig. 2D).

**Fig. 2.**
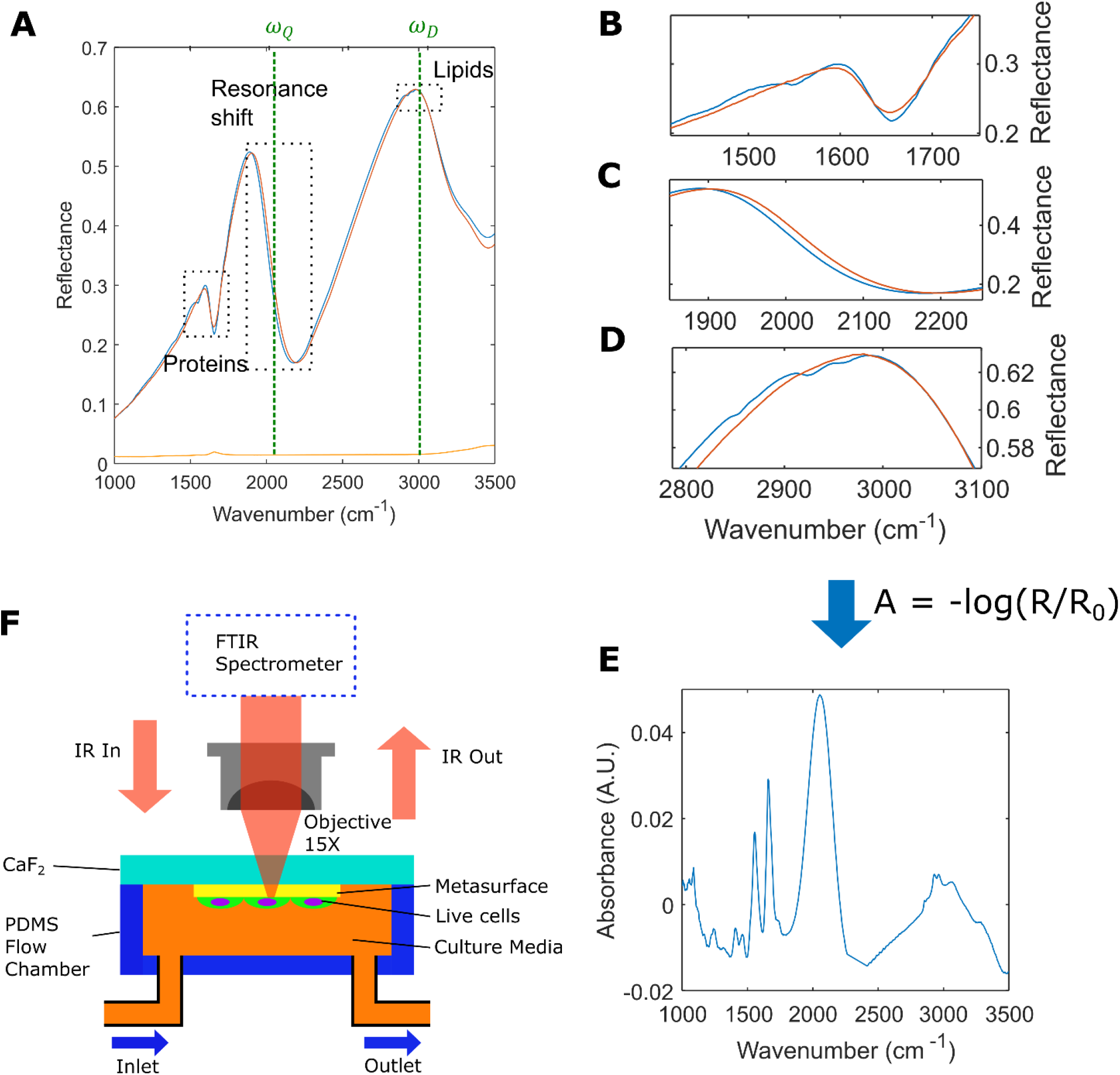
Measuring live cells with metasurface. **(A)** Reflectance spectra from the metasurfaces: *R*(*ω*) (with: blue line) and *R*_0_(*ω*) (without: red line) A431 cells. Reflectance spectrum from cells on a bare CaF_2_ substrate is shown in orange for comparison. Green dashed lines indicate the positions of dipolar (*ω*_*D*_) and quadrupolar (*ω*_*Q*_) resonances. Spectral regions showing the most difference between the two spectra are identified as **(B)** protein/amide absorption, **(C)** plasmonic resonance, and **(D)** lipids absorption windows. **(E)** Absorbance spectrum *A*(*ω*) = −log (*R*/*R*_0_) of A431 cells on the metasurface. The spectra are collected with unpolarized light. **(F)** Schematic drawing of the experimental setup.

These spectral features are presented in Fig. 2E as an absorbance spectrum *A*(*ω*) = -log (*R*/*R*_0_), where *R*(*ω*) and *R*_0_(*ω*) are the reflectance spectra with and without cells, respectively. The absorbance spectrum of A431 cells measured using MEIRS looks similar to the spectra collected from live cells using standard transmission or reflection techniques (*43, 44, 46, 58*), except with the addition of a strong Fano resonance shift peak around *ω*_*Q*_ ≈ 2080 cm^-1^. For brevity, we refer henceforth to the Fano feature as the *plasmonic resonance*. Because of the strong field concentration at the plasmonic resonance, it is extremely sensitive to any refractive index perturbation, regardless of whether its origin is in the cell or in the surrounding medium. In addition to protein-related amide and lipid-related C-H vibrations, additional asymmetric vibrations and bends of the C-H group at 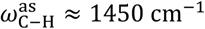 and 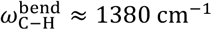 (lipids and proteins), as well as asymmetric phosphate stretch at 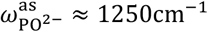 (phospholipids) are observed in the absorbance spectra. This confirms that the metasurface is sensitive to IR absorption as well as the refractive index changes originating from the cells.

### Cell response to low-concentration trypsin: intracellular signaling and detachment

Trypsin is a proteolytic enzyme commonly used to dissociate adherent cell culture from containers. We use trypsinization as a convenient method to validate MEIRS as a tool for real-time monitoring of cellular changes, and to characterize the overall cell-related IR signature. The perfusion medium was changed from L15 to Dulbecco’s phosphate buffered saline (DPBS), then trypsin in DPBS to cause cell dissociation. Dissociation of confluent layer of cells from the metasurface is continuously monitored until complete detachment of all cells (Fig. 3A). Preceding cell dissociation, we have observed two rapid effects described below: (i) refractive index modulation from the changing perfusion medium in the flow chamber (from L15 to DPBS), and (ii) activation of an intracellular signaling pathway by low-concentration (0.025%) trypsin solution in DPBS.

**Fig. 3.**
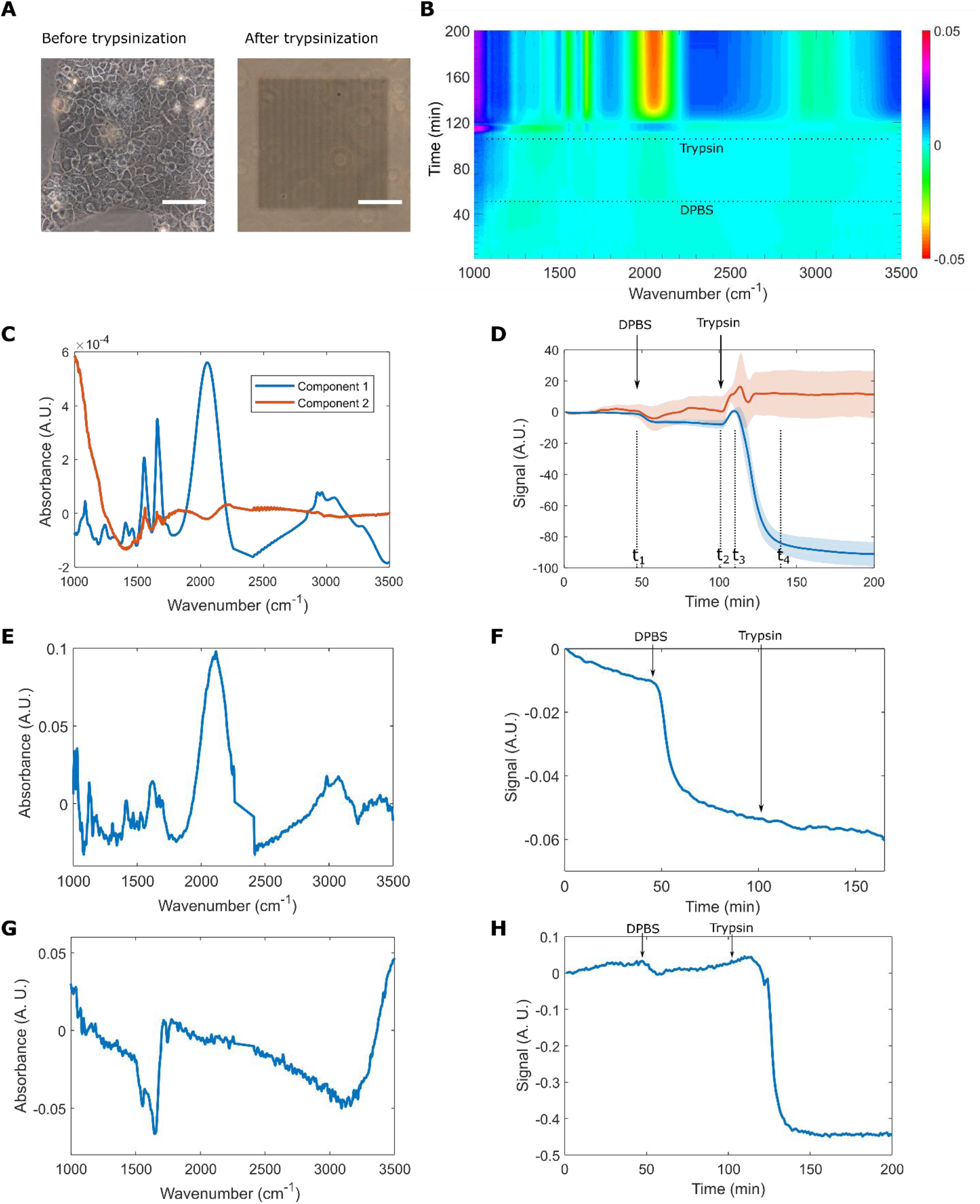
MEIRS monitoring of A431 cells trypsinization using sequentially injected DPBS and 0.025% trypsin-EDTA in DPBS. (**A**) Phase contrast microscopy images of the metasurface with cells before and after trypsinization. Scale bar: 100 µm. (**B**) A two-dimensional color plot of the phenotypic response barcode (PRB), showing the differential absorbance *A*(*ω, t*). Dotted lines indicate the timing of DPBS and trypsin injection. (**C**) Full spectrum component 1 (blue lines) and component 2 (red lines) spectral loadings 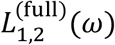. (**D**) Full spectrum time scores 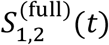. Solid line: the mean, shaded region: the standard error of the mean for a triplicate of experiments. Significant cellular response can be seen at *t*_1_ ≈ 45 mins, *t*_2_ ≈ 100 mins, *t*_3_ ≈ 110 mins, and *t*_4_ ≈ 140 mins. (**E**)-(**F**) Control experiment 1: metasurface but no cells. 1^st^ principal component spectral loading is shown in (E) and time score is shown in (F). (**G**)-(**H**) Control experiment 2: cells seeded on CaF_2_, no metasurface. 1^st^ principal component spectral loading is shown in (G) and time score is shown in (H). Arrows: the timing of DPBS and trypsin arrivals.

The reflectance spectrum *R*(*ω, t*) is continuously collected between *t*_0_ = 0 and *t*_fin_ = 200min while these stimuli are introduced. The resulting time-dependent differential absorbance spectra *A*(*ω, t*) = -log[*R*(*ω, t*)/*R*(*ω, t*_0_)] are presented as a two-dimensional PRB shown in Fig. 3B. Typically, multivariate analysis and classification techniques such as principal component analysis (PCA), partial least square (PLS) analysis, and hierarchical classification are used to analyze such spectral data (*46, 59*). In this work, we apply several dimensionality-reduction approaches based on PCA and factor rotation to understanding the timing and the underlying biochemical effects of the stimuli on live cells (see Methods for details).

First, PCA is used to analyze the entire spectral window, from 1000 cm^-1^ to 3500 cm^-1^. The two lowest spectral loadings 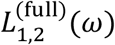 of the full-spectrum analysis are shown in Fig. 3C. Each experimental condition was repeated in triplicates, and the characteristic spectra obtained using PCA and factor rotation on one data set are then used as reference spectra for linear regression on subsequent experimental data sets to obtain the temporal scores 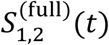 of the cells plotted in Fig. 3D (see Methods for details). The results of two control experiments, metasurface without the cells (Control 1: Fig. 3E,F) and cells on CaF_2_ without the metasurface (Control 2: Fig. 3G,H) have also been similarly analyzed for the same sequence of stimulants delivery.

Component 1 has a characteristic spectrum 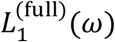 that includes all the major IR absorption peaks, including amides I and II, CH_2_/CH_3_ vibration, minor peaks in the fingerprinting region corresponding to carbohydrates and phospholipids, as well as a strong plasmonic resonance peak. We attribute this component to the presence/absence of cells on the metasurface, which directly corresponds to cellular adhesion. The temporal behavior of its corresponding time score 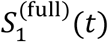 has three well-expressed features (see Fig. 3D). The small drop at *t* = *t*_1_ corresponding to the switching from L-15 medium to DPBS is attributed to refractive index modulation that leads to shift of the plasmonic resonance at *ω* = *ω*_*Q*_. This interpretation is backed by the prominence of the plasmonic feature in 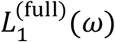. The increase in 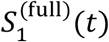 at *t* = *t*_2_, corresponding to the arrival of trypsin, is interpreted as being related to intra-cellular signaling (see below). Finally, the rapid reduction of 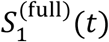 during the *t*_3_ < *t* < *t*_4_ time interval is attributed to cell detachment from the metasurface.

Note that the changes of the refractive index and composition of the injected fluid can also be observed in the Control 1 experiment, as shown in Figs. 3E,F. The spectral loading shown in Fig.3E features numerous vibrational lines corresponding to organic compounds, thus indicating the replacement of the nutrients-rich L15 medium by DPBS. The arrival of trypsin is not detectable in the Control 1 experiment because of its low concentration.

The enhancing role of the metasurface for cell monitoring is illustrated by comparing the temporal behavior of 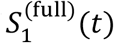 to its counterpart from the Control 2 experiment plotted in Fig. 3H. While the two are qualitatively similar for the DPBS arrival and cell detachment events, trypsin-induced intracellular signaling is unresolved without the metasurface. The origin of the spectral change without metasurface is likely to be from the displacement of cellular matter with water during cell dissociation from the substrate, as seen from the spectral loading peaked at the water absorption band (*ω* ∼ 3,500 cm^-1^ and *ω* ∼ 1,650 cm^-1^: see Fig. 3G). Unlike 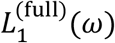, the control experiment spectrum (Fig. 3G) only shows a few weak vibrational lines that are attributable to the cells.

The interpretation of component 2 in Fig. 3C,D is less straightforward. Its spectral loading is characterized by a large change in the baseline level between 1000 cm^-1^ – 1400 cm^-1^, with a notable lack of any vibrational peaks in its spectrum 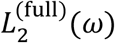. We conjecture that this spectral feature is due to Mie scattering from the cells (*60*), related to their rounding and morphological change as the cells dissociate from the metasurface and from each other. However, with the exception of the peak around *t* ≈ *t*_2_ seen in all replicates, there is a large variance between them in the corresponding time score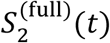, limiting its utility. Thus, the full-spectrum analysis of cell trypsinization yields only one highly-reproducible component, thereby reducing the phenotypic response to a single time-dependent score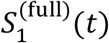. While different events are clearly detectable, the buffer-related refractive index change at *t* = *t*_1_ and the cellular response at *t* = *t*_2_ cannot be easily distinguished. Such one-dimensional cellular response is similar to those obtained from other all-optical analytic techniques (*47*–*53*).

To extract additional biochemical information, we selected three spectral windows: amides and lipids absorption regions, as well as the plasmonic resonance region, and performed the PCA on each of them separately. For simplicity, we focus only on the time-dependent scores of the first principal components for each spectral window. The corresponding spectral loadings 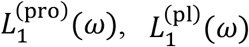, and 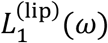, and their respective time scores 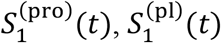, and 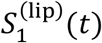, are shown in Fig. 4.

**Fig. 4.**
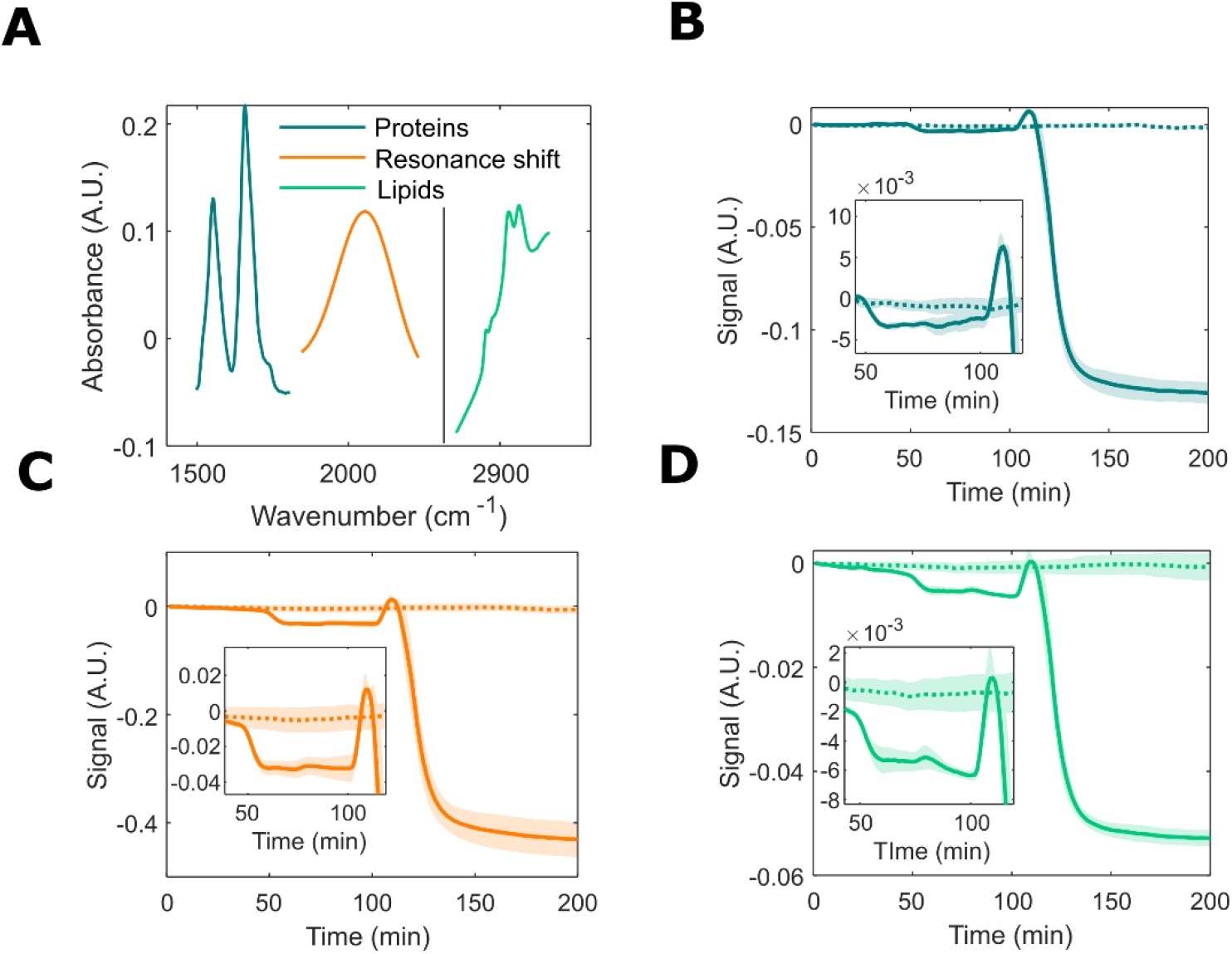
Analysis of trypsinization spectra using PCA over select spectral windows: proteins (1499 cm^-1^ < *ω* < 1807 cm^-1^), plasmonic (1845 cm^-1^ < *ω* < 2231 cm^-1^), and lipids (2756 cm^-1^ < *ω* < 3064 cm^-1^) absorption. The 1^st^ principal component is used for this analysis. (**A**) Characteristic spectral loadings in protein window (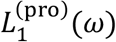, blue curve), plasmonic window (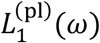, orange curve), and lipid window (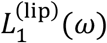, green curve), plottedinside their respective spectral windows. (**B**)-(**D**) Corresponding time scores: 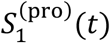 in (B), 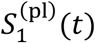in (C), and 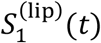 in (D), for the three spectral windows. Solid curves: DPBS (*t* ≈ 45 mins) and trypsin (*t*_2_ ≈ 100 mins) sequence as in Fig.3. Dotted curve: control with continuous perfusion of L-15 medium. Insets are enlarged to emphasize pre-detachment cellular response. Solid/dotted curves: the mean, shaded regions: the standard error of the mean for a triplicate of experiments.

The replacement of the L-15 media by DPBS at *t* ≈ *t*_1_ is clearly observed in all the time scores. Such change can be seen even for a bare metasurface without any cells (Control 1 experiment: dot-dashed line in Fig. S2), indicating that various amino acids, sugars, and other organic compounds in L-15 media could be partly responsible for this change. However, with A431 cells, the changes in signal level associated with DPBS arrival is consistently larger, suggesting that at least some of this signal change can be attributed to cellular response. Such additional response could be a result of the calcium-dependent changes in cellular adhesion, or cell volume changes as a result of change in osmolarity (*61, 62*).

More importantly, it is now possible to distinguish between the two responses at *t* ≈ *t*_1_(perfusion medium exchange) and *t* ≈ *t*_2_ (intra-cellular signaling), which have different relative magnitudes for the three scores. Specifically, by comparing the respective changes 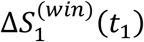and 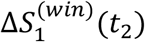 at the arrival times of DPBS and trypsin (here *win* stands for each of the three narrow spectral windows: protein, lipid, and plasmonic resonances), we conclude that the two responses at *t* ≈ *t*_1_ and *t* ≈ *t*_2_ are of entirely different nature: 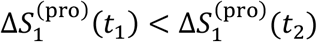 in the proteins window, but 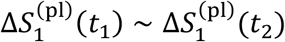 and 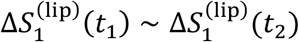 in the two remaining spectral windows.

Interestingly, for *t*_2_< *t* < *t*_3_, the sign of the cellular response 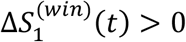 is positive in all three spectral windows, suggesting an increase in cellular signal, even though trypsin is known to cause cell dissociation. Such rapid cellular response induced by trypsin has been previously observed by RWG as a positive DMR (*63*). It has been attributed to the trypsin-induced activation of protease-activated receptors (PARs) belonging to a family of G protein-coupled receptors (GPCRs), which are endogenously expressed in A431 cells (*64, 65*). The agreement in the observed signal between MEIRS and RWG sensors confirms that at least part of the signal seen by MEIRS reflects changes in cellular adhesion, cytoskeletal reorganization, and GPCR-related signaling. Moreover, that the largest 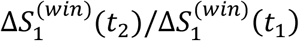 ratio corresponds to the protein window confirms that the mechanistic reason for trypsin-induced positive DMR is intracellular protein transport to the membrane caused by signaling (*65*).

### Cholesterol depletion in cellular membrane

The trypsinization measurement demonstrates the utility of MEIRS for measuring live cells. Next, we take advantage of the extreme concentration of the MIR optical field within a narrow penetration depth, which has been proven to be beneficial for analyzing protein monolayers and lipid bilayers (*17, 18, 20, 57*), to measure changes in cell membrane. For this, we applied MEIRS to detect the response of A431 cells to membrane cholesterol depletion induced by methyl-β- cyclodextrin (MβCD).

Cholesterol is a major component of the cell membrane in mammalian cells, influencing membrane fluidity and elasticity (*66*–*69*). MβCD, a water-soluble oligosaccharide with a hydrophobic cavity, is a compound widely used to manipulate membrane cholesterol. When applied to live cells, MβCD extracts cholesterol from the cellular membrane without entering the cells. Conversely, MβCD-cholesterol complexes (MβCD-chol) can be used to enrich membrane cholesterol. Cholesterol depletion through MβCD is known to trigger several different cellular responses, including ligand-independent activation of epidermal growth factor receptor (EGFR) (*66, 67*) as well as cytoskeletal reorganization that leads to higher cortical tension, increased focal adhesion size, and decreased cell spreading area (*68, 69*).

In our experiment, 10 mM of either MβCD or MβCD-chol dissolved in L-15 media were injected into the flow chamber, and the IR spectrum was continuously monitored. As with the trypsinization experiment, we first used the full-spectrum PCA to reduce the complexity of the data and identify the major features of cellular response (Figs. 5A and 5B). Components 1 and 2 have spectral signatures similar to those from trypsinization, and are attributed to cellular detachment from metasurface and Mie scattering from cell morphology change, respectively. Previously reported (*68, 69*) morphological changes of cells that eventually result in rounding and cell detachment have been confirmed by optical microscopy (Fig. S3). Here, we find that the cellular response can be roughly divided into three phases marked in Fig.5B and described below.

**Fig. 5.**
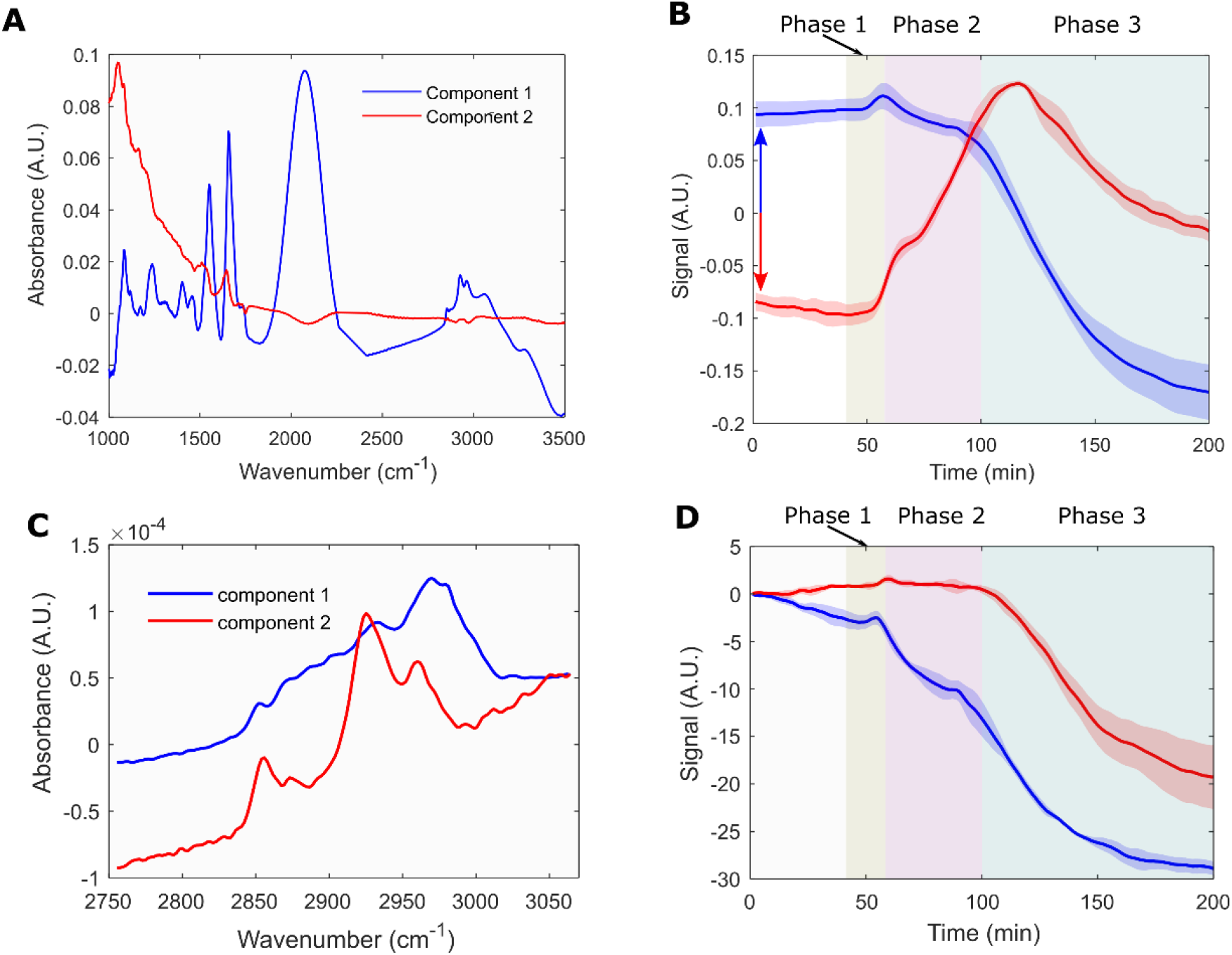
MEIRS of cholesterol depletion from plasma membranes of A431 cells by MβCD. (**A**) Full-spectrum spectral loadings 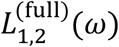 and (**B**) time scores 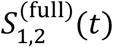. Component 1 (blue curve) is attributed to cellular adhesion, while component 2 (red curve) is attributed to change in cell morphology. Three phases of cellular response are shaded in the background to guide the eye. Red and blue arrows indicate that the temporal response curves are plotted with offset for clarity of presentation. (**C**) Lipid window spectral loadings 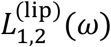 and (**D**) time scores 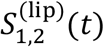. Component 1 (blue curve) is attributed mainly to cholesterol, while component 2 (red curve) is attributed to other cellular lipid contents. Note that component 1 shows sharp decrease upon the introduction of MβCD (Phase 2), whereas component 2 only starts decreasing once the cells start to detach (Phase 3).

Phase 1, which starts immediately after MβCD reaches the flow cell (*t*_1_ = 45 min) and lasts until *t*_2_= 55 min, is characterized by an increase in the score 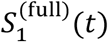of the first component. This increase is primarily caused by the refractive index increase due to MβCD dissolved in the media, as confirmed in a parallel experiment with no cells, see Fig. S4. Phase 2 starts at *t* = *t*_2_ and lasts until *t*_3_ = 100 min; it is characterized by a rapid decrease in *S* (*t*), followed by a plateau after 10 – 20 mins. During the same time, 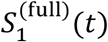 steadily increases, reflecting significant changes in cell morphology during this phase. Unlike component 2 in trypsinization experiments (see Fig. 3D), 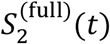 is very reproducible. Finally, Phase 3 starts at *t* = *t*_3_ and lasts until all cells are detached from the metasurface; both 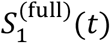 and 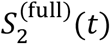 decrease during this phase.

A key question is whether it is possible to identify cholesterol depletion from the cellular membrane using two inherent capabilities of MEIRS: extreme optical field localization and spectral resolution. To this end, PCA was performed on the earlier defined lipids absorption region to extract the fine details of the spectral loadings 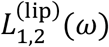 shown in Fig. 5C. Crucially, the above three phases of cellular response are clearly observed even when we analyze just the lipid absorption region, as seen from the time scores 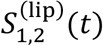 shown in Fig. 5D. Specifically, 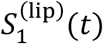 significantly decreases during Phase 2 of the cellular response, whereas 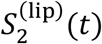 only startsdecreasing during Phase 3. To interpret these results, we note that the spectral signature 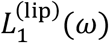 is similar to the absorption spectrum of cholesterol in the lipids region (Fig. S5): it is very broad, with few distinguishing features except for a peak at *ω* ≈ 2970 cm^-1^. The ability of high-concentration MβCD to affect other membrane lipids, including phospholipids, sphingolipids and other sterols, may explain this additional peak (*31, 58*). In contrast, 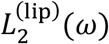 is characterized by three distinct absorbance peaks at 2854 cm^-1^, 2925 cm^-1^, and 2960 cm^-1^, which are attributed to the lipid acyl chains. Although precise peak assignment is challenging, the spectral change in the lipid spectrum during Phase 2 – captured primarily by component 1 – is different from that during Phase 3, during which the lipid absorption decreases mainly due to cell detachment. From this, we can conclude that indeed MEIRS can detect the rapid change in membrane lipid caused by MβCD during phase 2, and that, according to 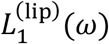, it is most likely related to cholesterol depletion.

Next, we compare the response of A431 cells to MβCD and MβCD-chol (Fig. 6) by analyzing the spectra inside the three earlier defined spectral windows. While all three phases show different time-dependencies of the spectra in response to the two compounds, the final Phase 3 is the easiest to interpret. In Phase 3, MβCD treated cells start detaching from the metasurface, and a decrease in IR absorbance score 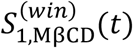 is observed in all three spectral regions. No comparable signal decrease in 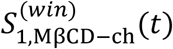 is found for any of the spectral windows because no cell detachment occurs in response to MβCD-chol.

**Fig. 6.**
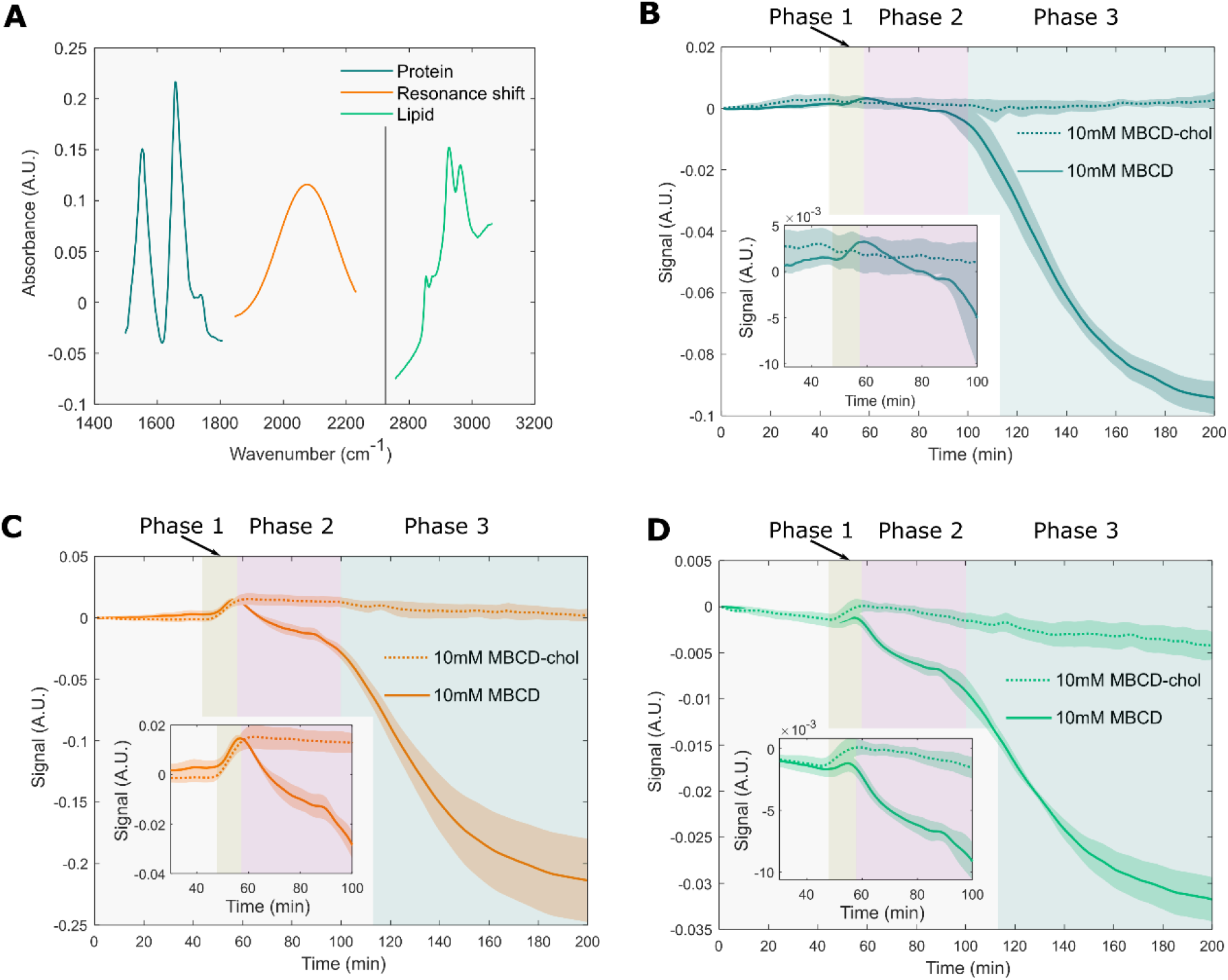
MEIRS of cellular response to MβCD and MβCD-chol using PCA in three spectral windows. The 1^st^ principal component is used for this analysis. (**A**) Characteristic spectral loadings in protein window (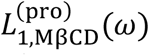, blue curve), plasmonic window (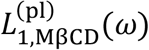, orange curve), and lipid window (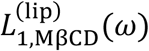, green curve), plotted inside their respective spectral windows. (**B**)-(**D**) Corresponding time scores: 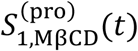 and 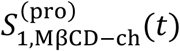 in (B), 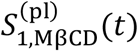 and 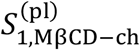 in (C), 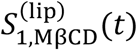 and 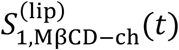 in (D). Solid (dotted) curves: 10 mM MβCD (MβCD-chol) in L-15 medium. Insets are enlarged to emphasize pre-detachment cellular response. MβCD/ MβCD-chol is injected into the flow cell at *t* ≈ 45 min. Three phases of cellular response are shaded in the background to guide the eye. Note that the lipids and plasmonic signals change almost immediately upon the introduction of MβCD, whereas the protein component starts changing only after the cells start to detach (Phase 3). Solid/dotted curves: the mean, shaded regions: the standard error of the mean for a triplicate of experiments.

During Phase 1, MβCD and MβCD-chol produce approximately equal refractive index changes, resulting in 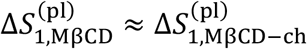. Also, the respective time scores 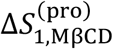and 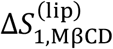 are small in the protein and lipid spectral windows. This implies that the observed signal does not originate from cells, but rather from changes in refractive index of the medium. This observation underlines an advantage of MEIRS over optical techniques that produce one-dimensional cellular signal such as SPR and RWG: these techniques would not be able to establish the origin of this signal, and to distinguish between optical signals produced by the cells and perturbations in the environment.

In contrast to Phase 1, spectral change from MβCD treatment occurring during Phase 2 is mainly attributed to cholesterol extraction from cellular membrane, which reduces the lipid absorbance and the local refractive index at the metasurface: see the respective decreases in 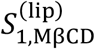 (Fig. 6D) and 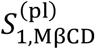 (Fig. 6C). Notably, the protein signal 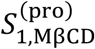 shows little change during Phase 2, suggesting that the adhesion sites formed by the cells on the metasurface remain intact. For comparison, MβCD-chol treatment results in little signal change after Phase 1, suggesting that cholesterol enrichment produces minimal phenotypical response from the cells. The different spectral response shown in the three spectral regions demonstrate that MEIRS is a powerful technique with detection specificity towards different families of compounds.

### Spectro-chemical imaging of live cells

IR spectro-chemical imaging of live cells is a powerful tool for analyzing biological samples such as tissue biopsies and fixed/live cells (*31, 32, 58*). For example, small quantities of highly heterogeneous cancer cells collected using minimally invasive biopsies benefit from imaging with cell-level imaging because drug-resistant minority of cells can be detected in a mixed population (*70*–*72*). Here, we demonstrate the suitability of MEIRS to spectroscopic imaging of sparsely seeded live A431 cells cultured on a metasurface. The spatially-resolved absorbance spectral hypercube *A*(*ω*, ***r***) = -log[*R*(*ω*, ***r***)] was collected from an array of imaged pixels located at ***r*** = (*x, y*) by coupling a focal plane array (FPA) with an FTIR-microscope system. The spectro-chemical image was obtained from *A*(*ω*, ***r***) using PCA to extract the spectral components originating from the cells (see Methods for the definitions of spectral loadings *L*_*i*_(*ω*) and spatial scores *S*_*i*_(***r***) used as spectral images).

There is some uncertainty as to whether metasurfaces are suitable for imaging, since some fabrication error is inevitable, and this may affect the plasmonic resonance and hence the image quality. Indeed, we found that the 1^st^ principal component (32% of total variance) corresponds to a gradual spatial variation in the plasmonic resonance (see Fig. S6 for the plots of *L*_1-3_(*ω*) and *S*_1-3_(***r***)). On the other hand, the 2^nd^ principal component (PC2) spectral loading *L*_2_(*ω*) (12% of total variance) shown in Fig. 7B exhibits high similarity to the spectral change observed from cell detachment due to trypsinization shown in Fig. 3C (component 1). This agreement strongly suggests that PC2 corresponds to the IR absorbance from the cells. Indeed, the IR spectro-chemical image corresponding to *S*_2_(***r***) shown in Fig. 7A matches well with the visible microscopy image of the same cells shown in Fig. 7C,D.

**Fig. 7.**
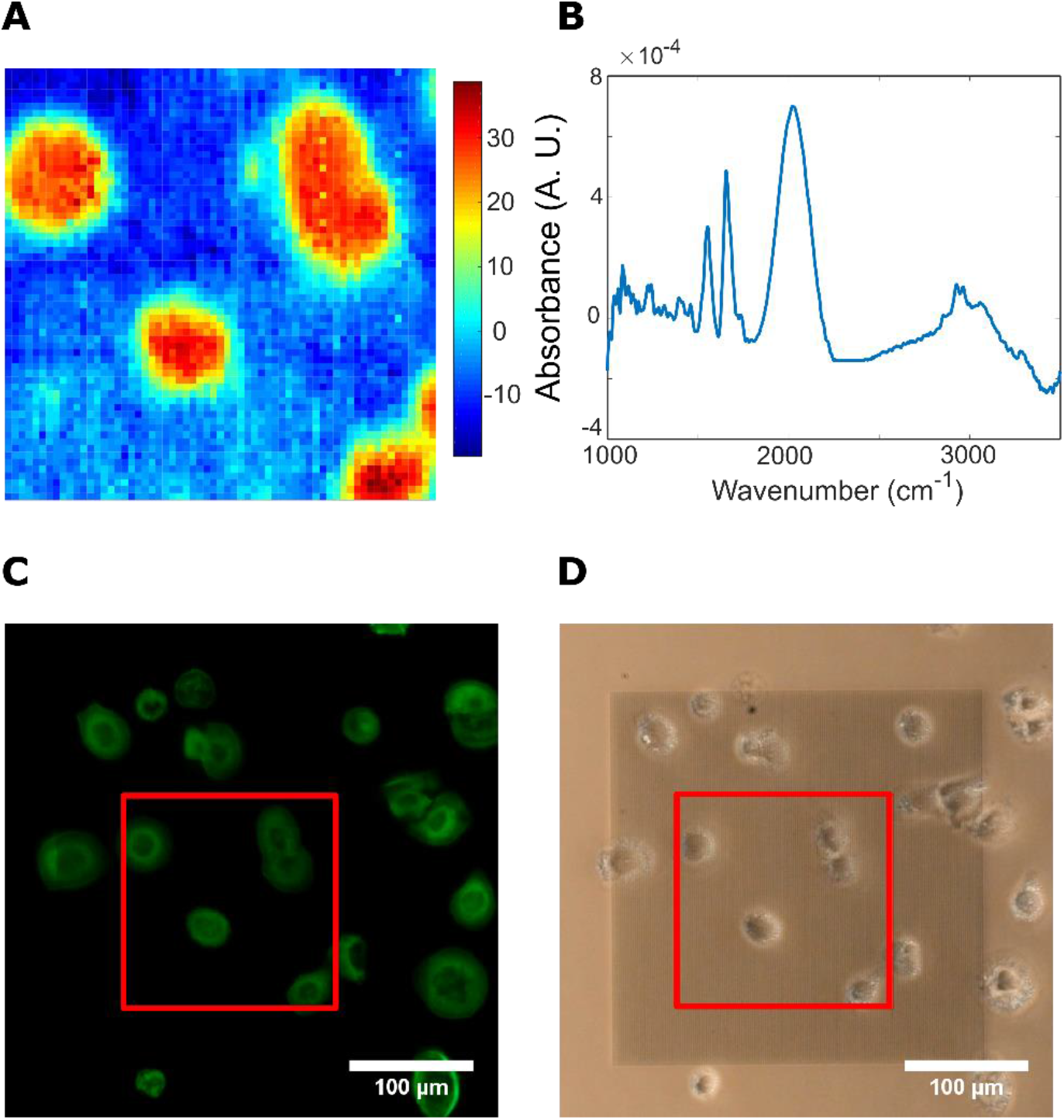
Imaging live cells on metasurface using FPA. (**A**) Spectro-chemical image *S*_2_(*r*) of A431 cells on the metasurface corresponding to the score of 2^nd^ PC (12% of the variance). Imaged region corresponds to a 173 µm × 173 µm square region. (**B**) Corresponding spectral loading *L*_2_(*ω*) associated with the image in (A). (**C**) Fluorescence and (**D**) phase contrast image of the cells imaged in (A). Red squares: the region imaged using FPA. Cells are stained using phalloidin, showing the cytoskeletal structure. Scale bar: 100 µm.

Although each individual cell can be readily identified, the finer sub-cellular features are not resolved. While we do not expect to see the organelles and nucleus that are deep in cell cytosol due to the limited penetration depth of the metasurface, typically there is also significant variation in the cell attachment on the basal side, with focal adhesions in close contact with the substrate and larger gaps between the cell and the substrate in the other areas (*73*). The absence of such variation from the spectro-chemical image *S*_2_(***r***) is attributed to the diffraction-limited lateral spatial resolution, which is approximately 5 µm–15 µm resolution in the wavenumber range of 3000 cm^-1^ –1000 cm^-1^. Nevertheless, the single-cell imaging with MEIRS opens new and exciting opportunities for label-free detection of the response of individual cells to various external stimuli in a heterogeneous mixture. Because of the extreme sensitivity of MEIRS to cell adhesion, it may be possible to detect adhesion changes much earlier that the overall morphological changes detected by phase imaging.

## Discussion

We have demonstrated the application of MEIRS – a metasurface-based nearfield IR spectroscopic assay – to the real-time, label-free characterization of live cells and their responses. MEIRS can be used to detect and distinguish cellular responses to chemical stimuli from the distinct spectroscopic signatures associated with each of them. The detected cellular responses, encoded into a two-dimensional phenotypic response barcode (PRB), vary from cell adhesion and morphological changes to intra-cellular signaling and protein transport. The near-field nature of MEIRS enables us to study subtle changes in the molecular composition of cellular membranes, such as cholesterol depletion. In this work, we have analyzed PRBs using a relatively simple multivariate PCA technique. The use of more advanced machine learning models such as deep neural networks may further assist with classifying various sequential and concurrent cellular processes.

In combination with an FPA imaging system, MEIRS was also shown to be a promising non-perturbing technique for spectro-chemical imaging of live cells with single-cell resolution. Although sub-cellular features cannot be resolved because of diffraction limit, imaging cells using MEIRS can be useful for the identification of different sub-populations of cells in a heterogeneous population (e.g., primary cells from biopsy) and for the detection of their distinct responses to therapeutics.

In this work, we have found that A431 cells preferentially adhered to the gold nanoantennas, rather than the CaF_2_ substrate. In general, cells may interact with nanostructures in a complex manner, but a thorough investigation of such interaction is beyond the scope of this work. Engineering surface chemistry and nano-topography of the metasurface may further expand MEIRS by allowing it to study effects related to cell-nanostructure interaction, such as membrane deformation and endocytosis, which are interesting topics for future studies.

MEIRS has a promising application as a label-free cellular assay for drug screening. Our metasurface technology has much similarity in device geometry to well-established phenotypical cellular assays based on electrical impedance and resonant waveguide grating. As we have demonstrated in this work, MEIRS is fundamentally superior to these technologies because, as a spectroscopic technique, it produces information-rich PRBs that enable us to better elucidate the underlying phenotypic responses. Multivariate analysis enables us to distinguish between different cellular responses, as well as to separate environmental perturbations from cellular signals. Although our current implementation is based on a fluidic cell, metasurfaces are fabricated on planar, transparent CaF_2_ substrates, which makes the technology scalable and allows for integration with microplate geometry. This would pave the way to a high-throughput IR spectroscopic assay that can probe cellular responses to various compounds, as well as their interaction with nanostructured surfaces.

## Materials and Methods

### Fabrication of metasurface

The Fano resonance plasmonic metasurfaces are fabricated on 12.5mm x 12.5 mm x 0.5mm CaF_2_ substrates using electron beam lithography, electron beam evaporation of gold, followed by lift-off. Polymethyl methacrylate (PMMA) e-beam resist is spin-coated on the CaF_2_ substrate, followed by another layer of DisCharge to reduce electron charging. The metasurface pattern is defined using e-beam lithography with JEOL 9500 system. The exposed resist is developed using 1:3 MIBK:IPA developer. Gold metasurface are deposited using electron beam evaporation of 5 nm of Cr, followed by 70 nm of Au, and the samples are lift-off in acetone overnight.

### Cell culture

A431 human epidermoid carcinoma cell line (acquired from ATCC) is used as a model system. Cells with passage number < 15 are used for all experiments. The cells are cultured in Dulbecco’s modified Eagle medium (DMEM) supplemented by GlutaMAX, 10 % fetal bovine serum (FBS), and 1% penicillin/streptomycin in a standard incubator with 5 % CO_2_ and 37 °C. Prior to seeding the cells on the metasurface, the metasurface is treated with 10 µg/mL fibronectin in phosphate buffered saline for 1h at 37 °C. The metasurface is then placed in a 12-well plate, and cells are seeded in DMEM at approximately 200,000 cells per mL. The cells are then allowed to proliferate until confluent. Prior to a measurement, the cells on metasurface are serum-starved overnight.

### IR spectroscopy

For IR spectroscopy, the metasurface with A431 cells is attached to a PDMS flow cell, with a total volume of approximately 20 µL. The IR spectra of the metasurface are measured using an FTIR spectrometer (Bruker Vertex) coupled to an IR microscope (Bruker Hyperion 3000), fitted with a reflective Cassegrain objective and a mercury-cadmium-telluride (MCT) detector. The flow cell is perfused with L15 media supplemented with 1% antibiotic-antimycotic at a flow rate of 0.1 µL/s. A microscope stage heater is used to maintain the flow chamber at 37°C. The measurement is made in reflectance mode, with the IR light going through CaF_2_ substrate. Unpolarized light is used for the measurement. FTIR spectra are collected at 1 acquisition/minute, 120 averaging for both background and sample, at 4 cm^-1^ spectral resolution. Mertz phase correction and 3-term Blackman-Harris apodization function are used.

### Trypsinization measurement

The flow cell is perfused with L15 medium at a flow rate of 0.1 µL/s for 1h prior to the measurement using a programmable syringe pump. L15 medium is chosen to maintain physiological pH level under ambient atmospheric condition.

The following sequence of stimuli injection is introduced. First, Dulbecco’s phosphate-buffered saline (DPBS) with no calcium and magnesium is injected into the chamber at *t*_1_ ≈ 45 mins. Next, a 0.025% (10μM) trypsin-ethylenediaminetetraacetic acid (EDTA) solution in DPBS is injected into the chamber at *t*_2_ ≈ 100 mins. Cell detachment starts approximately at *t*_3_ ≈ 110 mins and completes at approximately *t*_4_ ≈ 140 mins. The reflection spectrum *R*(*ω, t*) is continuously collected at a sampling rate of *v* = 1 spectrum/min over the *t*_0_ < *t* < *t*_fin_ ≡ 200 mins time interval. Each experiment is repeated in triplicate with separate cell culture.

### Cholesterol depletion measurement

The flow chamber is first perfused with L15 medium at a flow rate of 0.1 µL/s for 1h prior to the measurement. 10 mM MβCD or 10 mM MβCD-chol solution in L-15 media is injected into the flow cell at approximately 45 mins at 0.1 µL/s. Each experiment is repeated in triplicate with separate cell culture.

To prepare MβCD-chol, 10 mM MβCD solution in L-15 medium is mixed with excess cholesterol and agitated overnight at 37 °C. The excess cholesterol is filtered out with 0.2 µm syringe filter prior to using the solution.

### FTIR data analysis

Water vapor spectra are subtracted from the sample spectra using an in-house MATLAB code. Spectra between 2262 cm^-1^ and 2412 cm^-1^ are cut out to remove the contribution from CO_2_ line. The reflectance spectra are vector normalized and converted to absorbance spectra. The spectra are then smoothed using 11 point Savitzky–Golay filter.

The entire PRB data set consists of *N* = *N* × *M* (where *N* = *vt*_fin_ = 200 time points and *M* = 1303 spectral points) data points, which is standardized and analyzed using PCA. We have carried out the full-spectrum PCA using *N* multi-dimensional vectors *A*(*ω, t*) collected at different time instances as distinct data points. Such analysis, equivalent to a rotation in the *M*-dimensional space, establishes a set of principal component (PC) loadings *L*_*i*_(*ω*) sorted in such a way that *L*_1,2…*M*_(*ω*) account for progressively smaller spectral variance. Typically, just a few *m* ≪ *M* PCs are needed to account for over 95% of the temporal evolution of *A*(*ω, t*), constituting a major dimensionality reduction of the data set. Therefore, by neglecting all principal components with *i* > *m*, we can approximate the evolving spectra as

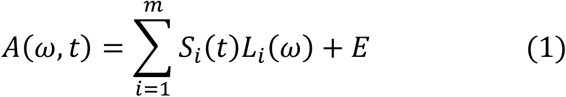

where *S*_*i*_(*t*) is the time-dependent score of the i’th PC with the spectral loading *L*_*i*_(*ω*), and E is an error matrix. The significance of the PC loadings is that they can be interpreted as corresponding to specific biochemical characteristics evolving with time, e.g., overall lipid content or composition. Their corresponding time scores indicate how these characteristics evolve with time. An additional promax rotation in the reduced *m*-dimensional space is used to improve the interpretability of the spectra. For brevity, we use the same notation (*S*_*i*_(*t*) and *L*_*i*_(*ω*)) for the rotated component time scores and spectral loadings.

PCA is performed for both the full spectral region, or smaller spectral regions corresponding to the absorption peaks of proteins, lipid, as well as plasmonic shift. For PCA of amides, plasmonic resonance shift, and lipids spectral regions, the spectral windows of 1499 cm^-1^ – 1807 cm^-1^, 1845 cm^-1^ – 2231 cm^-1^, and 2756 cm^-1^ – 3064 cm^-1^, respectively, are used. PCA is used to extract the characteristic spectra using one set of experimental data out of triplicates. The PCA loadings are then used as reference spectra in linear regression to obtain the temporal scores for the remaining data sets, using the regression model

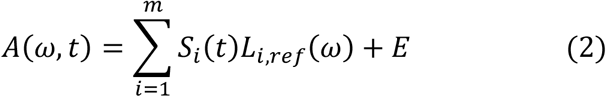

where *L*_*i,ref*_(*ω*) are the reference spectra obtained from PCA loadings.

### FPA imaging

FPA imaging is performed using an FPA detector coupled to Hyperion 3000 IR microscope. 15X, 0.4 NA reflective Cassegrain objective is used with a 64 × 64 pixel FPA, with a final pixel size of 2.7 µm. Spectra are acquired at 4 cm^-1^ resolution and 120 averaging, with each sample scan taking approximately 10 min. The metasurface is integrated with the flow cell, with the cells kept under continuous perfusion of L-15 medium throughout the measurement as described earlier. For simplicity, the cells are not subjected to any chemical stimuli during imaging.

Fabrication imperfection in the metasurface results in spatial non-uniformity in the resonance spectrum of plasmonic antennas across the metasurface. Because each A431 cell covers many plasmonic antennas, it is difficult to distinguish between the natural complexity of the cell shape and unintentional non-uniformity of the antennas. We utilize PCA to separate these two spectral changes.

We apply PCA to absorbance defined as *A*(*ω*, ***r***) = -log[*R*(*ω*, ***r***)], where ***r*** = (*x, y*) is the location of the imaged pixel. The result of PCA can be expressed as 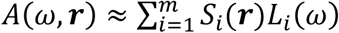, where *S*_*i*_(***r***) is the space-dependent score of the i’th principal component with the spectral loading *L*_*i*_(*ω*). If the i’th spectral loading corresponds to IR absorption by the cell, then *S*_*i*_(***r***) can be interpreted as corresponding to a spectro-chemical image of a single cell, or multiple cells shown in Figs.7 C,D. On the other hand, those spectral loading that do not contain any expected vibrational fingerprints can be dismissed as originating from unintentional nonuniformity of plasmonic antennas or noise.

### Fluorescence staining

A431 cells grown on the metasurface are fixed with 4% paraformaldehyde solution for 15 minutes, followed by permeabilization using 0.5% Triton X-100 for 10 minutes. The sample is then washed with PBS and stained using Alexa Fluor 488® phalloidin for F-actin labelling.

### Numerical simulation

The near-field profile and field enhancement of the metasurface are simulated using COMSOL Multiphysics, a commercially available software using the finite elements method. The geometric parameters are taken from the SEM measurement in Fig. 1A, with a gold thickness of 70 nm. The metasurface array is simulated in unit cell with Floquet boundary condition in x and y direction. As in the experiment, the metasurface is simulated with water on the top and supported by CaF_2_ on the bottom, both layers with thickness equal to a least one wavelength. The wave excitation is simulated using a period port, where y-polarized plane waves are incident normally from CaF_2_ side. A Perfectly Matching Layer boundary condition is used at the end of the water side.

## Author Contributions

SHH and GS conceived and designed the experiment. SHH performed the experiments. JL and ZF performed the numerical simulations. SHH, JL, and RD fabricated the metasurface. SHH and GS analyzed and interpreted the data. GS supervised the project. The manuscript was written through contributions from all authors.

## Acknowledgements

The research reported here was supported in part by the National Cancer Institute of the National Institutes of Health under award number R21 CA251052 and in part by the National Institute of General Medical Sciences of the National Institutes of Health under award number R21 GM138947. This work was performed in part at the Cornell NanoScale Facility, a member of the National Nanotechnology Coordinated Infrastructure (NNCI), which is supported by the National Science Foundation (Grant NNCI-1542081). This work also made use of the Cornell Center for Materials Research Shared Facilities which are supported through the NSF MRSEC program (DMR-1719875).

