## Supplementary Information for "Monitoring the effects of chemical stimuli on live cells with metasurface-enhanced infrared reflection spectroscopy"

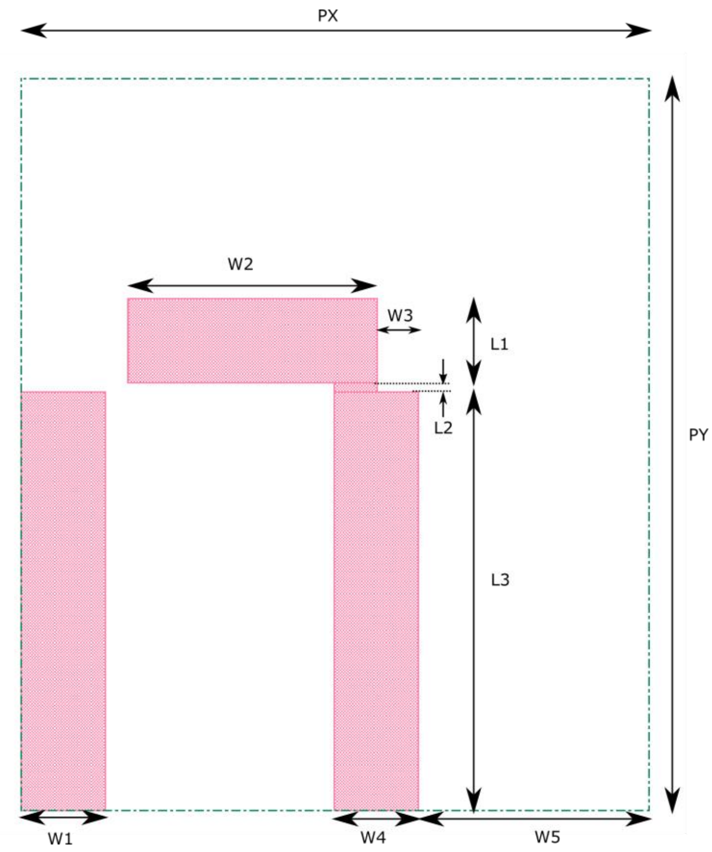


**Fig. S1.**

**Detailed geometry of the metasurface.** W1 = 194 nm, W2 = 578 nm, W3 = 97 nm, W4 = 194 nm, W5 = 535 nm, L1 = 194 nm, L2 = 22 nm, L3 = 972 nm, PX = 1458 nm, PY = 1701 nm.

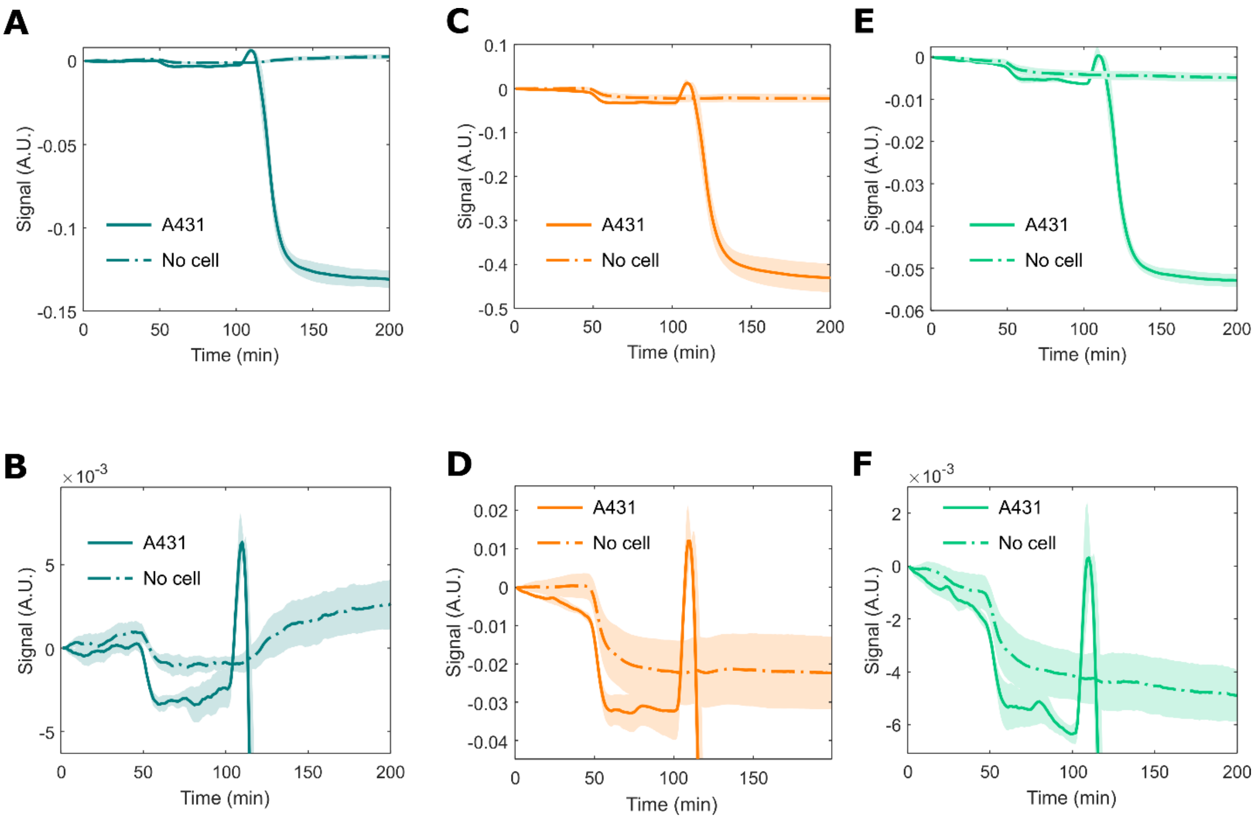


**Fig. S2.**

**Comparison between metasurface with A431 cells and metasurface with no cells, in response to DPBS and trypsin injection.** DPBS arrives at the flow cell at around 45 min, while trypsin arrives at 100 min. Solid curve: time score in each spectral region for metasurface with A431 cells (same data as presented in Fig. 4). Dotted curve: time score in each spectral region for a bare metasurface. (**A**)-(**B**) Protein region. (**C**)-(**D**) plasmonic region. (**E**)-(**F**) lipid region. (B), (D), (F) show the same data as in (A), (C), (E), but scaled to show the response from DPBS. Solid/dotted curve: the mean, shaded region: the standard error of the mean for a triplicate of experiments. Note that the exchange from L15 medium to DPBS results in spectral changes in all three spectral regions, with or without cells on a metasurface. However, metasurface with cells results in consistently larger spectral changes.

A

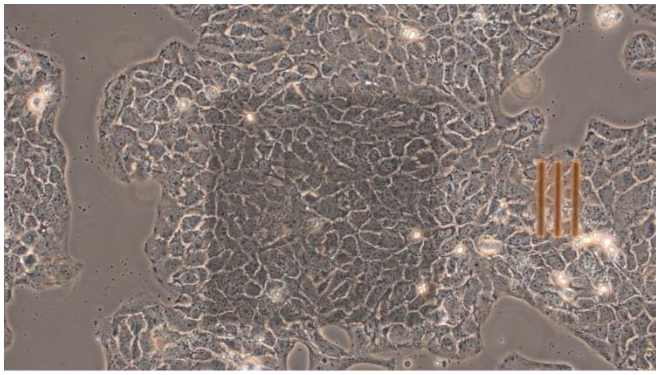

B

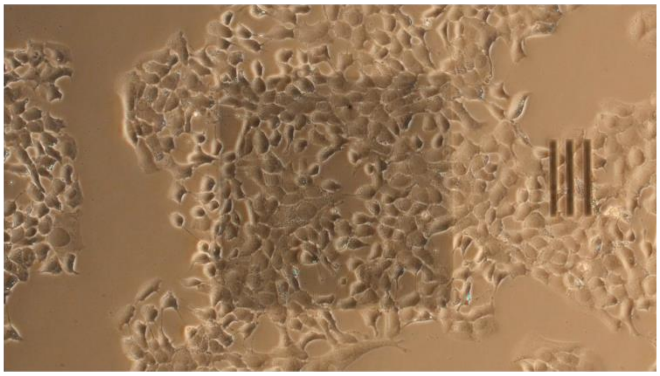

C

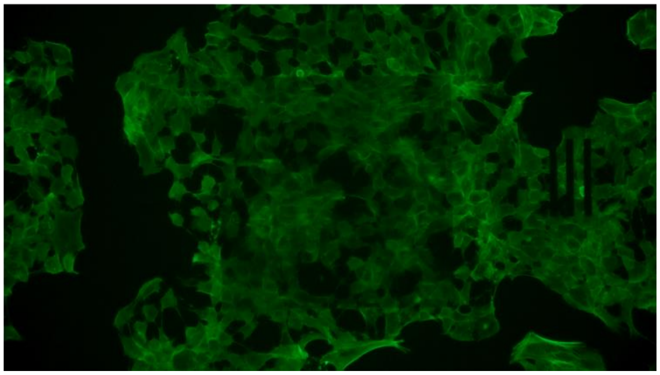

**Fig. S3.**

**Morphology change in A431 cells after treatment by 10 mM MβCD.** (**A**) Phase contrast image, before MβCD treatment. (**B**) Phase contrast image, 1h after MβCD treatment. (**C**) Phalloidin-stained fluorescence image, 1h after MβCD treatment. As can be seen from (B) and (C), upon MβCD treatment, cells begin to round, decreasing in area in contact with the metasurface. Tethering between cells can be observed as the cells are rounded. Eventually, cells are detached from the metasurface.

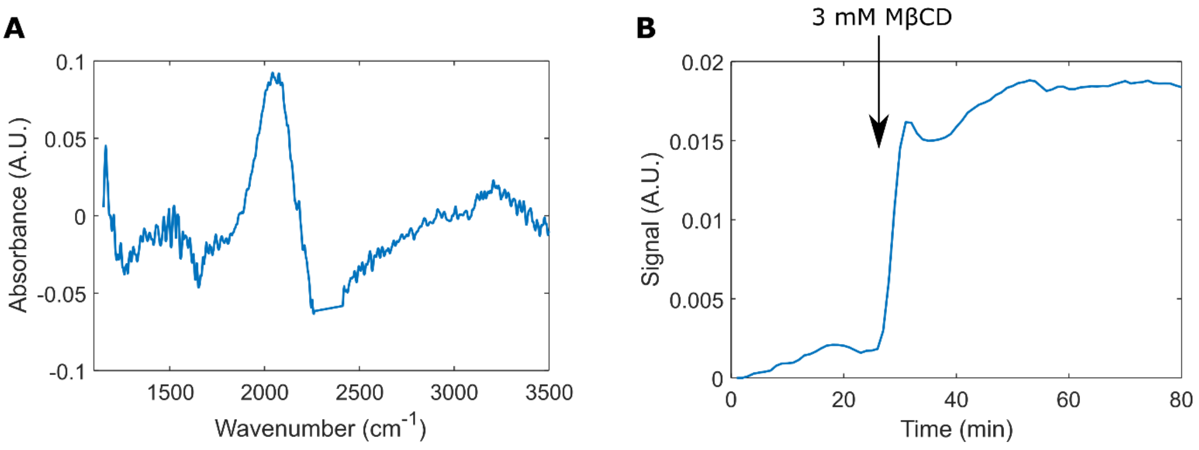


**Fig. S4.**

**Observed spectral changes on the metasurface with no cells, in response to 3 mM MβCD.** (**A**) 1st principal component spectral loading. (**B**) 1st principal component time score. The spectral signature of MβCD is dominated by a resonance shift, suggesting that refractive index change is the dominant change as seen by the metasurface.

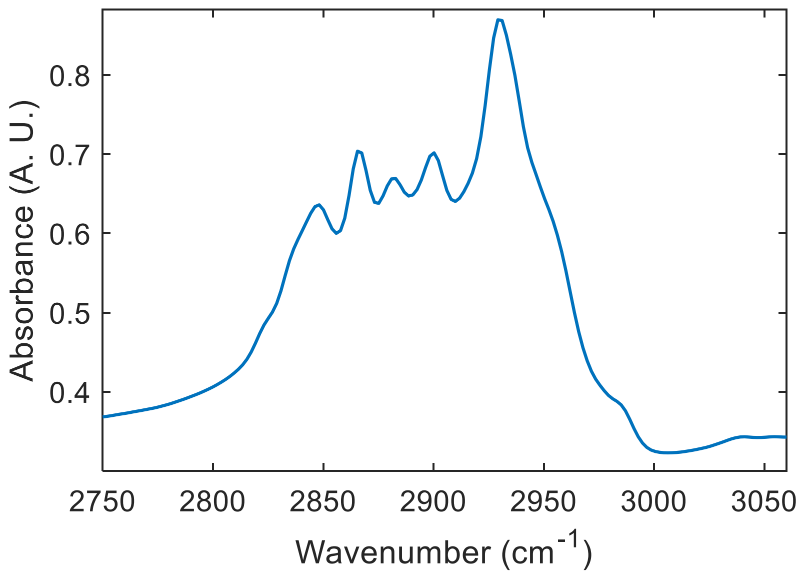


**Fig. S5.**
**Cholesterol absorption spectrum in the $CH_2/CH_3$ stretching region.**

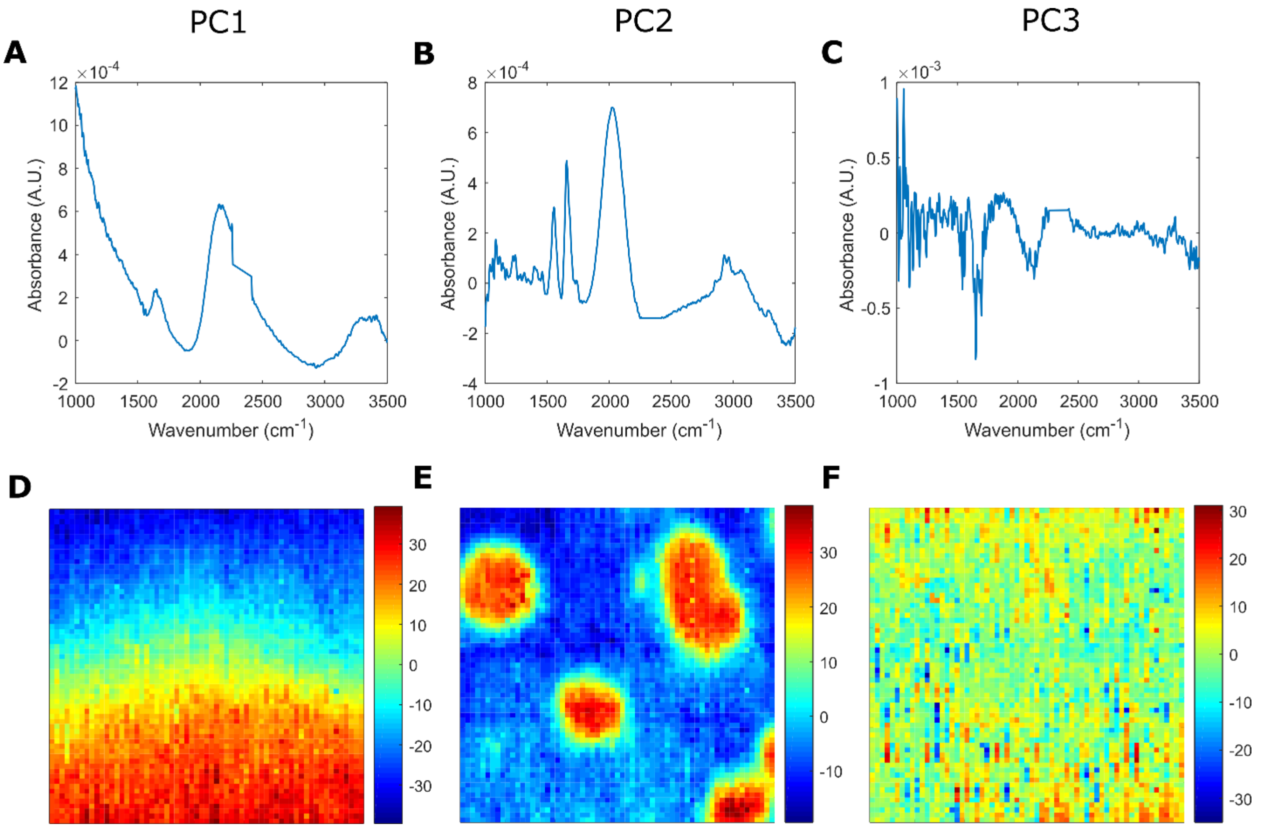


**Fig. S6.**

**Spectrochemical imaging using MEIRS.** First 3 principal components are shown. (**A**)-(**C**) Loadings of the first 3 principal components. (**D**)-(**F**) Score mapping of the first 3 principal components. PC1, PC2, and PC3 explain 32%, 12%, and 3% of the variance, respectively. The 1$^{st}$ principal component appears to correspond to slow variation in the baseline level, which is likely the result of small variation in the metasurface resonance across the measured pixel. The 2$^{nd}$ principal component corresponds to the signal from cells. The 3$^{rd}$ principal component and subsequent components appear to be dominated by noise.
